# Improved Yield for the Enzymatic Synthesis of Radiolabeled Nicotinamide Adenine Dinucleotide

**DOI:** 10.1101/2022.12.12.520150

**Authors:** Jared Eller, Shivansh Goyal, Xiaolu A. Cambronne

## Abstract

Labeled β-nicotinamide adenine dinucleotide (NAD) analogs have been critical for uncovering new biochemical connections and quantitating enzymatic activity. They function as tracers for enzymology, flux analyses, and in situ measurements. Nevertheless, there is limited availability of specific types of analogs, especially radiolabeled NAD analogs. Here, we describe an improved enzymatic synthesis reaction for ^32^P-NAD^+^ with a yield of 98% ± 1%, using lowered concentrations of reactants and standard equipment. This represents the highest reported yield for the enzymatic synthesis of NAD^+^ to date. With the high yield we were able to directly use the reaction product to generate derivatives, such as ^32^P-NADP. The high-yield enzymatic synthesis is versatile for a broad variety of labels and NAD derivatives. Its advantages include lowered concentrations of reactants, providing sufficient amounts of product for downstream applications, and minimizing intermediate purification steps.

---

β-Nicotinamide adenine dinucleotides (NAD) are conserved co-enzymes used by cells to facilitate reversible oxidoreductive reactions via transfer of hydride ions. In this scenario, the relative levels of oxidized dinucleotide (NAD^+^, NADP) compared to their reduced counterparts (NADH, NADPH) will influence the capacity for either oxidative or reductive reactions to occur. Distinctly, oxidized NAD^+^ and its derivatives (e.g. ADPR, cADPR, NAADP) have additional roles as signaling molecules and can regulate enzymatic activities as required co-substrates or ligands. In this second scenario, the concentration of available nucleotide can limit signal transduction. These distinct usages sometimes converge, and this has been shown as a mechanism to coordinate regulation of intracellular pathways during adipogenesis.^1-2^ Given the complexity of processes that depend on NAD metabolites, the concentrations of these molecules are tightly regulated inside and outside of cells, and its levels are compartmentalized subcellularly.^1,3^

Molecular analogs of NAD have been instrumental in advancing our understanding of this complex biology. They have been used to quantitate mass spectrometry measurements, to uncover regulation of flux, and to obtain enzymatic rates.^4-8^ By measuring specifically labeled NAD pools amid complex mixtures, researchers can quantitatively determine alterations in flux and concentrations. In the example of the mammalian mitochondrial NAD^+^ transporter^9-11^, kinetic rate measurements of NAD^+^ uptake in intact and respiring mitochondria were made feasible with radiolabeled NAD^+^, and investigation into the roles of mitochondrial NAD^+^ in cells was feasible with heavy-atom labeled NAD^+^.^9-12^ Labeled NAD^+^ molecules were distinguishable from the unlabeled resident pool, allowing small additions to be measured over the course of the experiments; this resolution could not have been achieved by simply measuring final mitochondrial NAD^+^ amounts. Moreover, specific advantages of radio- and heavy-labeled analogs include that they are structurally identical and recognized by endogenous enzymes as cognate substrates, and that they can be quantified with a variety of highly sensitive detection methods. Despite the broad utility of radiolabeled NAD analogs, the availability of these reagents is limited.

To address this, we sought to improve the yield of the enzymatic synthesis of NAD^+^ to facilitate the ability for individual labs to obtain specifically labeled NAD and NAD derivatives.^13-16^ We surmised that a higher yield reaction would be amenable for labeling experiments and for generating labeled NAD-derivatives. Moreover, we sought a high yield reaction that could be performed using standard equipment as a useful and versatile resource for many labs.

We considered an enzymatic strategy described by multiple groups, which relies on recombinant NMN adenylyl transferases (NMNAT enzymes) and ATP to generate β-NAD from either β-nicotinamide mononucleotide (NMN) or β-nicotinamide mononucleotide hydride (NMNH).^13-14^ This strategy had been adopted into a two-step synthesis to produce NAD^+^ derivatives such as nicotinamide 4’-thioribose NAD^+^ (S-NAD^+^) at 70% yield, starting from modified 4-thio nicotinamide riboside.^15^ We reasoned that the synthesis of labeled NAD analogs could be simplified into a single-step process by including radiolabeled analogs directly at the adenylyl transferase step (Figure 1a). We also wanted to test the consequence of including inorganic pyrophosphatase (PPase) in the reaction to degrade the inorganic pyrophosphate (iPP) byproduct, with the idea that this could drive the forward reaction for a higher final yield (Figure 1a). This idea was supported by the addition of PPase in the combined reaction that improved the yield of NAD^+^ by ∼10%.^16^ However, whether PPase improved the adenylation step was never directly tested, and if Mg^2+^ is ever limiting, a formal possibility is that PPase could cleave ATP and limit the reaction by depleting ATP instead of promoting the forward reaction. Thus, we directly tested whether the addition of PPase improved the adenylation reaction of NMN in vitro and whether this could be used to incorporate a ^32^P-label onto NAD^+^. An additional advantage is that the addition of PPase could eliminate the need for using initial reactants at high concentrations, which greatly facilitates the ability to use radiolabeled reactants. For comparison, the combined enzymatic synthesis reactions used millimolar concentrations of reactants.^15-16^ Notably, by simply adjusting the type of labeling on ATP or NMN, or using NMN derivatives (e.g. NMNH or nicotinic acid mononucleotide), this enzymatic synthesis could produce a variety of NAD-related nucleotides, highlighting the potential versatility of the approach. To monitor the reaction and to quantitate the yield, we used α-^32^P-ATP (3000Ci/mmol, 10 mCi/mL) that had a single label at its α-phosphate, thus quantitation of ^32^P in the product would directly correlate with product formation.

**Figure 1:**
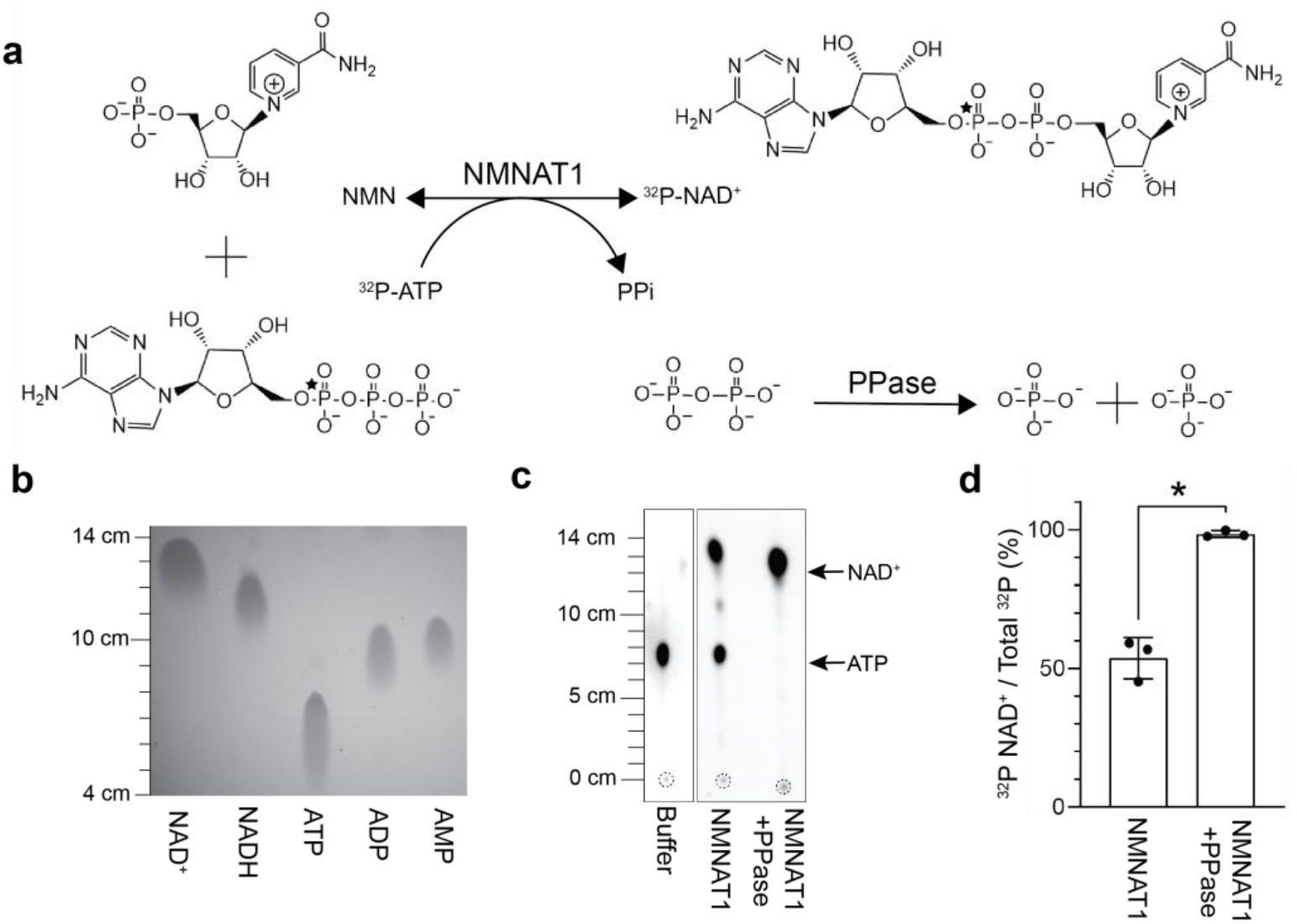
PPase improved the yield of the NMN adenylyl transferase reaction in vitro (a) Proposed reaction for the enzymatic synthesis of ^32^P-NAD^+^ from ^32^P-ATP. Position of the ^32^P-label is indicated by a star. (b) NAD^+^, NADH, ATP, ADP and AMP standards (>95% purity, 5 μL of 50 μM stocks) were resolved with thin layer chromatography (TLC) using PEI cellulose F plates and imaged following excitation at 254 nm. (c) 1 μL from the enzymatic reactions were resolved with TLC and quantitated following exposure with a phosphorimaging screen, representative images. Leading edges were measured from the origins (dashed circles). (d) Calculated yields of ^32^P-NAD^+^ (mean ± SD, n=3, Student’s t-test, *p< 0.01).

We used thin layer chromatography (TLC) to distinguish the synthesized products from its reactants.^17^ Briefly, cellulose plates were initially resolved to 5 cm above the origin using milliQ water in a lidded glass tank and air-dried. Plates were then resolved with 1.4 M LiCl to 15 cm above the origin.^17^ We quantitated the migration of known nucleotide standards and used this reference to identify ^32^P-labeled species in our reaction based on their migration patterns. To visualize the standards, we used PEI cellulose F plates with an incorporated fluorescent indicator for visualization of unlabeled molecules following 254 nm excitation (Figure 1b).

In parallel reactions, we combined 3 μM α-^32^P-ATP (≤ 120 μCi), 50 μM ATP, 50 μM NMN and 5 μM NMNAT1 in a final buffer of 50 mM Tris-HCl pH 7.4, 100 mM NaCl, 12 mM MgCl2 and 1 mM DTT. One unit of PPase was added to one of the reactions and omitted from the other. The reactions were incubated at 22 °C for 2 hours and 1 μL from each reaction was analyzed with TLC. To image the ^32^P-nucleotides, dried plates were wrapped in clear clingwrap and exposed to a phosphorimager screen for three minutes. Images were captured using a Typhoon scanner and individual spots were analyzed with ImageJ for distance of migration from origin and for their integrated intensities above background using the built-in Gel Analyzer feature. Integrated intensities for each spot were calculated as a percentage of the total intensity in the sample. The signal from ^32^P-NAD^+^ divided by the total ^32^P signal per lane was used to calculate the yield of the reaction.

We observed that addition of PPase significantly improved the yield of the enzymatic adenylation reaction. Without PPase, we observed a mean yield of 54% ± 7%, and substantial amounts of unconverted ^32^P-ATP, indicating that the reaction was at equilibrium (Figure 1c,d). This was consistent with previously reported yields of ∼70% in a combined reaction.^15^ In contrast, the addition of PPase appeared to prevent the main reaction from reaching equilibrium and resulted in a 98% ± 1% incorporation of the ^32^P signal into NAD^+^, corresponding to a yield of approximately 49 μM ^32^P-traced NAD^+^ (2.9 μM ^32^P-NAD^+^ and 46 μM unlabeled NAD^+^) (Figure 1c,d). Expected to be also remaining in the mixture would be ∼4 μM ^32^P-traced ATP (240 nM ^32^P-ATP and 3.8 μM unlabeled ATP), ∼1 μM NMN, as well as NMNAT1 and PPase enzymes. To stop the reaction, the tube was flash frozen with liquid nitrogen and stored at -20 °C.

In the reactions that did not include PPase, we observed ^32^P-reaction contaminants that migrated ∼10.5 cm and that only appeared when co-incubated with NMNAT1 (Figure 1c). Although this migration pattern corresponds to ^32^P-ADP/AMP it is unclear how these contaminants originated. One possibility is that the NMNAT1 preparation resulted in a slower side-reaction with the residual ^32^P-ATP. With the addition of PPase, appearance of the ^32^P-contaminants was lessened.

To confirm that we had synthesized ^32^P-NAD^+^, we used a freshly thawed fraction of the reaction product to test whether it could be imported by human SLC25A51, a mitochondrial transporter that preferentially imports oxidized NAD^+^ over reduced NADH, nicotinamide mononucleotide (NMN), and nicotinamide (Nam) at physiological concentrations.^9-10^ In previous work, we had showed that ectopic expression of human SLC25A51 complemented the selective import of NAD^+^ in isolated mitochondria from S. Cerevisiae that were genetically ablated for their ability to import NAD^+^ (DKO mitochondria, Δndt1 Δndt2 BY4727).^9^ In vitro, we observed significantly increased uptake of the ^32^P-labeled product by human SLC25A51-expressing DKO mitochondria compared to mitochondria expressing the empty vector control (Figure 2, Supporting information). This confirmed that ^32^P-NAD^+^ was the dominant, labeled species in the reaction. NMR analyses of parallel unlabeled reactions supported the conclusion that NAD^+^ was the dominant product and that addition of PPase promoted increased yields of NAD^+^ (Figure S1).

**Figure 2:**
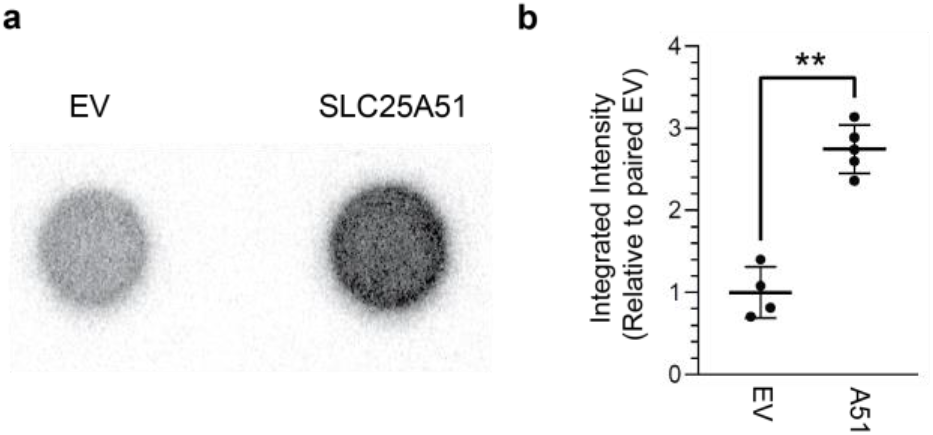
Selective uptake of radiolabeled reaction product by SLC25A51 (a) Representative ^32^P-dot blots of in vitro NAD^+^ import assays from DKO mitochondria that either expressed empty vector (EV) or a human mitochondrial NAD^+^ transporter (SLC25A51). (b) Relative mean intensities following uptake of ^32^P-product between paired EV and SLC25A51 conditions (mean ± SD, n=4-5, Student’s t-test, **p < 0.001).

The high yield adenylation reaction created an opportunity to directly use the product for generation of radio-labeled ^32^P-NAD derivatives. As proof-of-principle, we sought to generate ^32^P-NADP from ^32^P-NAD^+^ using recombinant human NAD Kinase (hNADK) and ATP (Figure 3a).^18-19^ In a 20 μL reaction, we combined 3 μL of the reaction product (∼440 nM ^32^P-NAD^+^ final, ≤18 μCi), 50 μM NAD^+^, 100 μM ATP, 10 mM Tris-HCl pH 8.0, 100 mM NaCl, and 5 mM MgCl2 with 5 μM hNADK. The reaction was incubated at 22 °C for 16 hours, and 1 μL of the reaction was analyzed by TLC. The generated reference image (Figure 3b) indicated that NADP migrates farther than NAD^+^ under the described TLC conditions. We observed that approximately 22% and 55% of the ^32^P-NAD^+^ is converted to ^32^P-NADP within two and eighteen hours, respectively (Figure 3c). The data provided a proof-of-principle that the high yield adenylation reaction generated by addition of PPase resulted in a sufficiently uniform α-^32^P-product for the direct synthesis of labeled derivative α-^32^P-NADP with a minimization of side-reactions.

**Figure 3:**
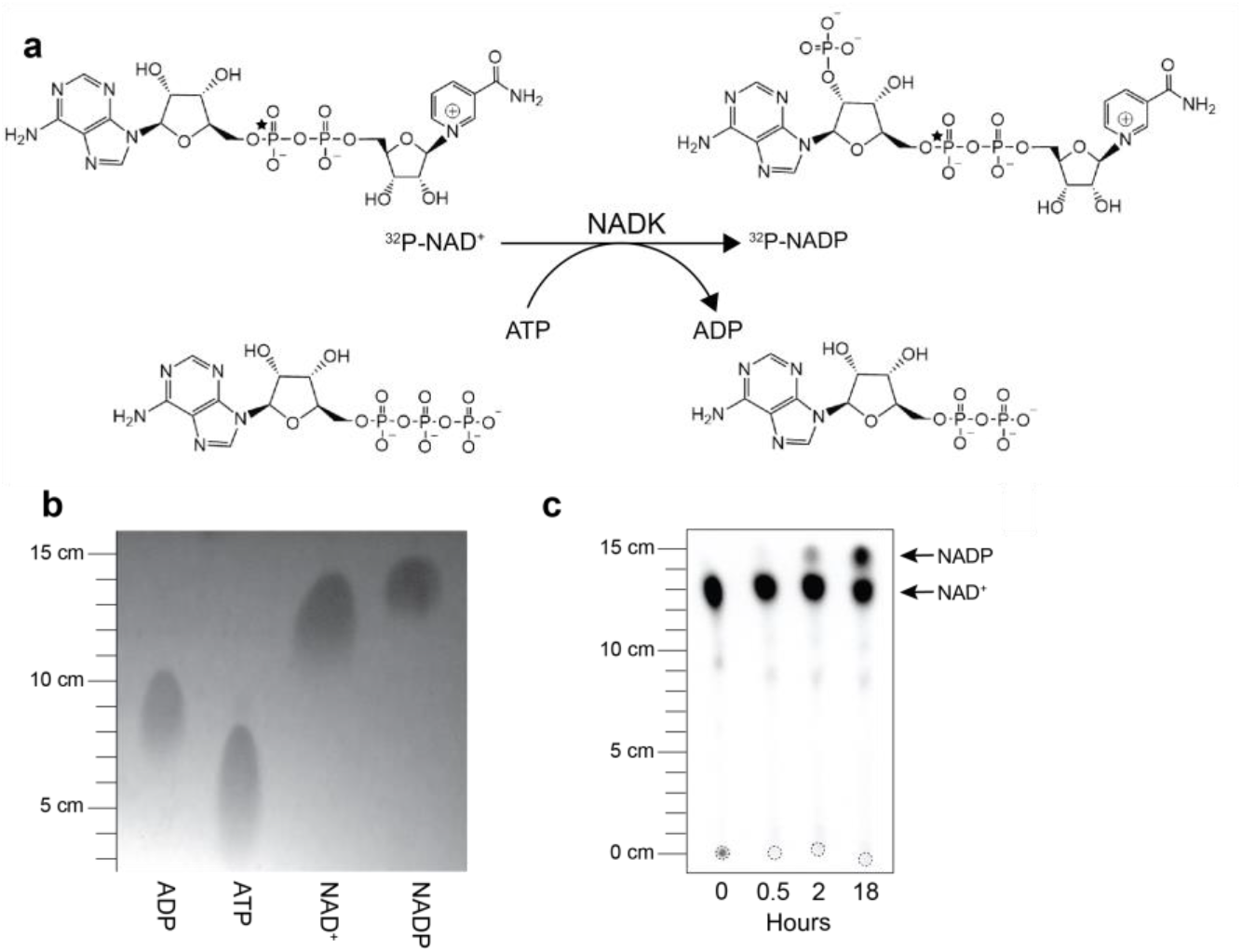
Direct synthesis of labeled NADP from the 32P-NAD+ reaction product. (a) Proposed reaction for the synthesis of labeled NADP from ^32^P-NAD^+^. Position of the ^32^P-label is indicated by a star. (b) ADP, ATP, NAD^+^, and NADP standards were resolved with TLC using PEI cellulose F plates and imaged following excitation at 254 nm. (c) Samples (1 μL) from the NADK enzymatic reaction were obtained at indicated time points, resolved with TLC, and radioactivity was detected with phosphorimaging. Leading edges were measured from respective origins (dashed circles).

In cases where a higher purity radiolabeled product is required, any resolvable ^32^P-traced nucleotide can be recovered following TLC. To demonstrate the feasibility of this technique, we performed an enzymatic synthesis without PPase and observed the expected product mixture including ^32^P-NAD^+^ and unreacted ^32^P-ATP (Figure S2a). We used a clean razor blade to isolate the cellulose from the TLC plate corresponding to ^32^P-traced NAD^+^ and extracted the nucleotide by filtration (Supporting Information). The recovered material was analyzed by TLC and predominantly contain the desired ^32^P-NAD^+^ (Figure S2a). This material could then be used in SLC25A51-dependent uptake assays, as described above (Figure S2b).

In conclusion, we have shown that addition of PPase in the NMN enzymatic adenylation reaction resulted in high-yield generation of NAD^+^, and this could be an effective method for generating labeled analogs. We expect that this approach could be expandable to work with other types of labels and placed in different positions. This high yield reaction required lowered concentrations of reactants compared to previously published approaches.^13-16^ Thus, there is opportunity for a broad range of reactants and more diverse set of possible products. Additionally, the labeled NAD^+^ product can be directly used to generate labeled derivatives with an additional enzymatic reaction, such as NAADP, NADP(H), NADH or ADPR, without further processing. Methods for creating NADH by alcohol dehydrogenase and ADPR by porcine brain NAD^+^ nucleosidase are described in Feldmann et al. ^16^ NAADP can be produced from NADP in vitro using CD38 at an acidic pH and high concentrations of NA.^20^ Notably the addition of PPase may not improve every adenylation reaction, highlighting the need to have empirically confirmed this approach. When attempting to test PPase with FAD synthetase (FADS) to recreate ^32^P-FAD from FMN and ^32^P-ATP, we observed that addition of PPase created an aberrant ^32^P product that migrated to ∼10.5 cm, and could represent AMP, ADP or cyclic-FMN (Figure S3).^21^ The appearance of a PPase-dependent aberrant product indicated that its utility may depend on the specific reaction.

Together, we have tested and described an approach that is versatile and easily adaptable for the improved enzymatic synthesis of labeled nicotinamide dinucleotides and derivatives. It has the potential to be expanded to include many other analogs and adenylated products with specific labels.

## ASSOCIATED CONTENT

### Supporting Information

This file contains Supporting Figures S1-S3, corresponding figure captions, a detailed experimental methods section, and an extended reference section for the Supporting Information.

## AUTHOR INFORMATION

### Author Contributions

JME, SG, and XAC conceptualized the project and contributed to experimental design. JME performed all experiments. The manuscript was written with contributions from all authors, and all authors have given approval to the final version of the manuscript.

### Accession Codes

NMNAT1, Q9HAN9; PPase, P0A7A9; NADK, O95544; FADS, Q8NFF5

### Notes

The authors declare no competing financial interests.

Any additional relevant notes should be placed here.

## ACKNOWLEDGMENTS

We thank Prof. Marie Migaud at the University of South Alabama for guiding us to consider the NMNAT1 enzymatic reaction, advice and discussion. We thank Prof. Adrian Keatinge-Clay and Dr. Takeshi Miyazawa at the University of Texas at Austin for guidance performing and analyzing the NMR experiments. Research in the laboratory was supported by funding from the National Institutes of Health (DP2 GM126897), the PEW Charitable Trusts, and Cancer Prevention and Research Institute of Texas (RP210079).

---

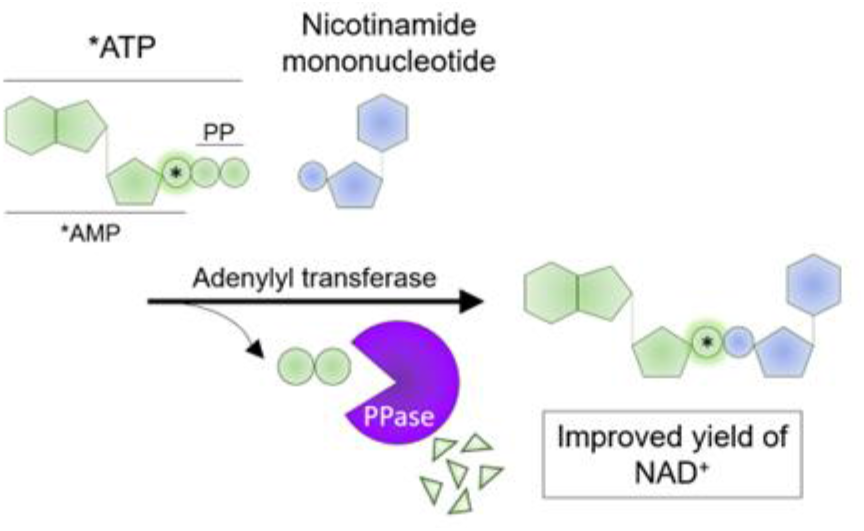

## Supporting Information

**Supporting Figure S1:**
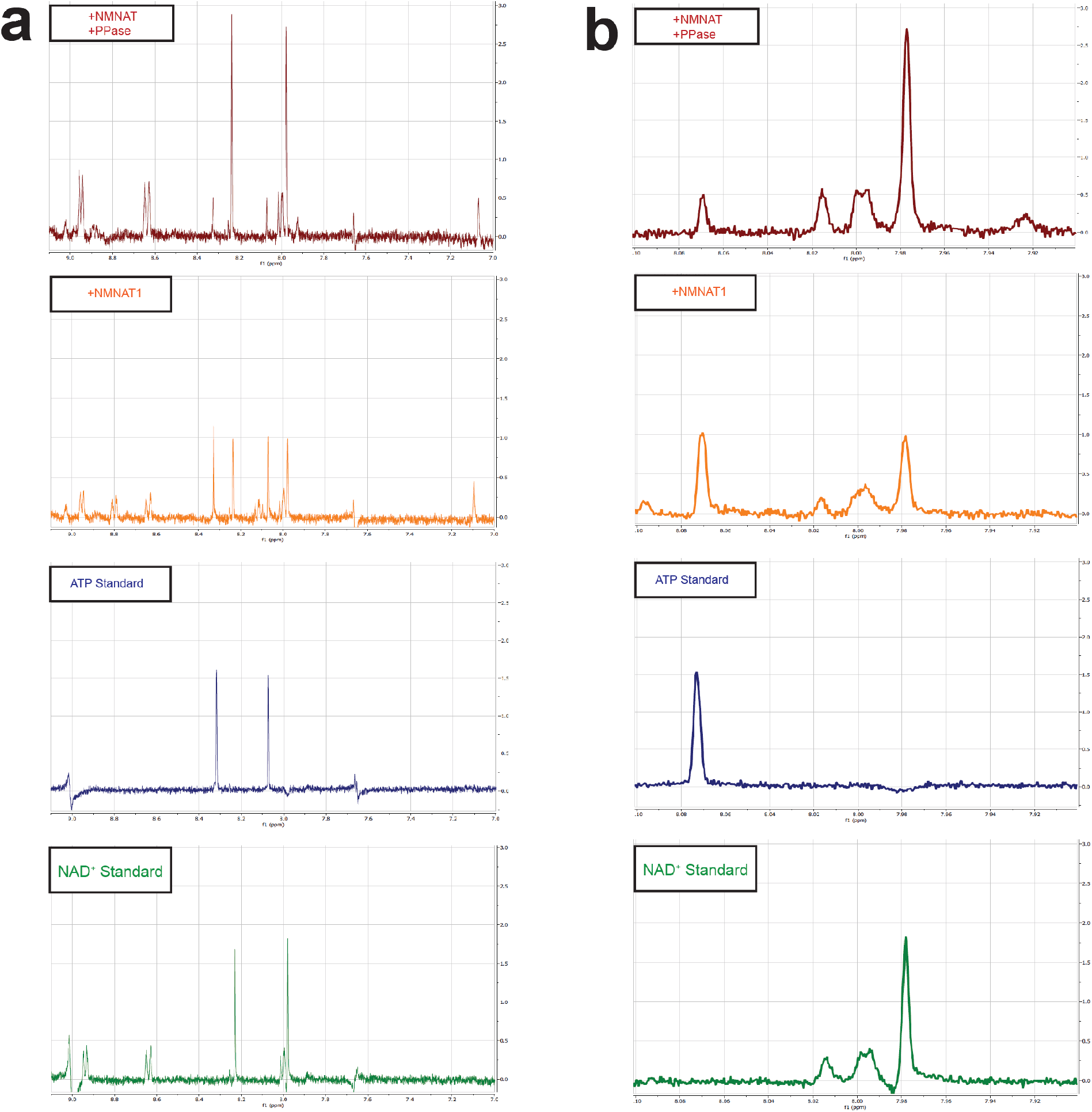
^1^H-NMR of NMNAT1 Reaction with and without PPase. Aligned ^1^H-NMR spectra of 1 mM standards for NAD^+^ (green) and ATP (purple) with comparable x- and y-axes. Panel (a) represents a broad view, 7.00 - 9.10 ppm. Panel (b) represents a focused view. 7.90 – 8.10 ppm, to better visualize a representative subset of peaks. In orange the reaction only contained NMNAT1 without PPase and it showed peaks of comparable areas characteristic of NAD^+^ and ATP peaks as determined by the standards. Maroon represents the combined reaction with NMNAT1 and PPase. Here we observed increased areas corresponding to NAD^+^ peaks and decreased areas corresponding to ATP peaks.

**Supporting Figure S2:**
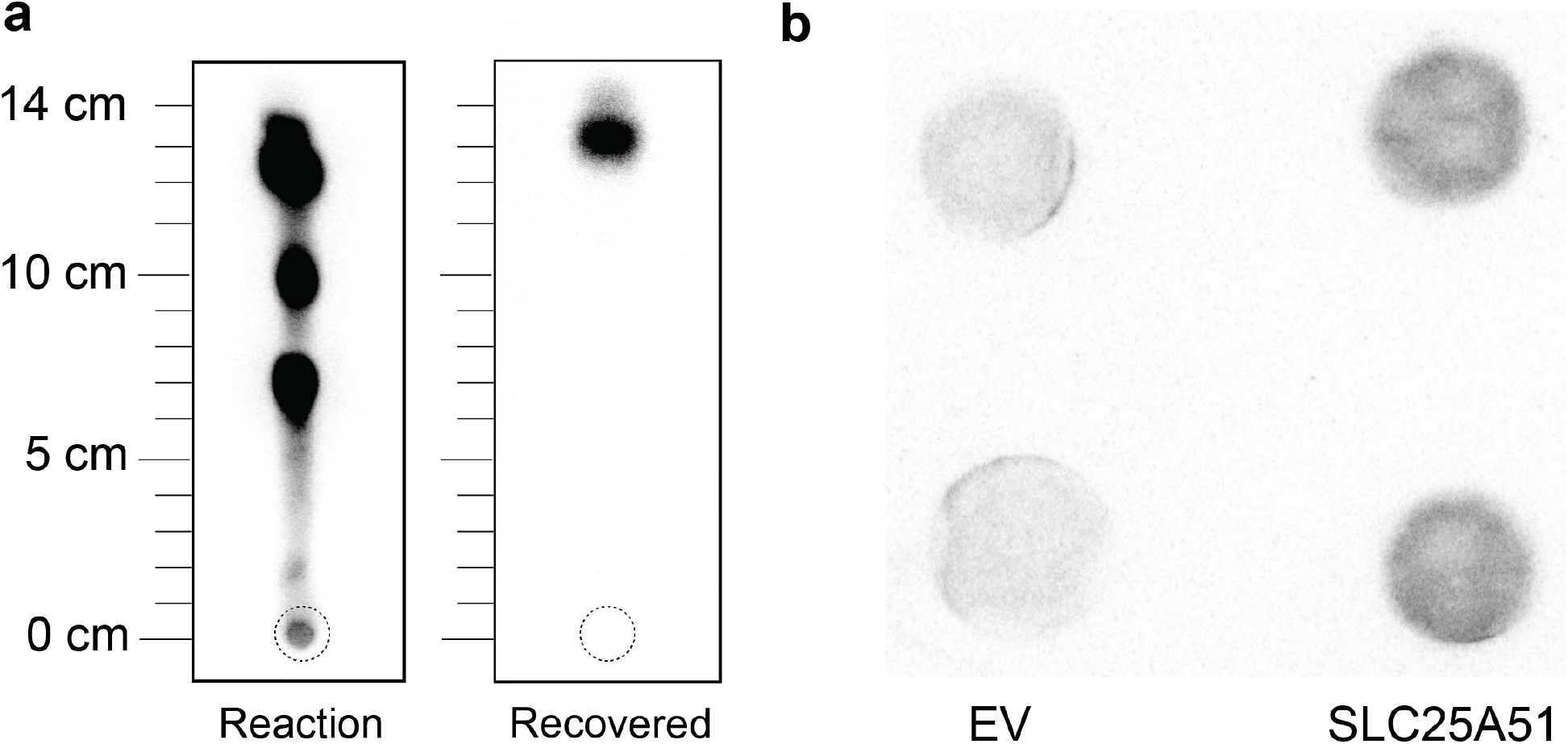
NAD^+^ Uptake with Recovered ^32^P-NAD^+^. (a) Thin layer chromatography (TLC) of the enzymatic NMNAT1 reaction in the absence of PPase show as expected a mixture of ^32^P-NAD^+, 32^P-ATP, and other ^32^P-species *(Reaction, left)*. 2.5% of total material from the recovered ^32^P-NAD^+^ sample was resolved using TLC (*Recovered, right*). (b) ^32^P-dot blots of in vitro NAD^+^ import assays using recovered ^32^P-NAD^+^ material and isolated yeast double knockout mitochondria that either expressed empty vector (EV) or human mitochondrial NAD^+^ transporter (SLC25A51).

**Supporting Figure S3:**
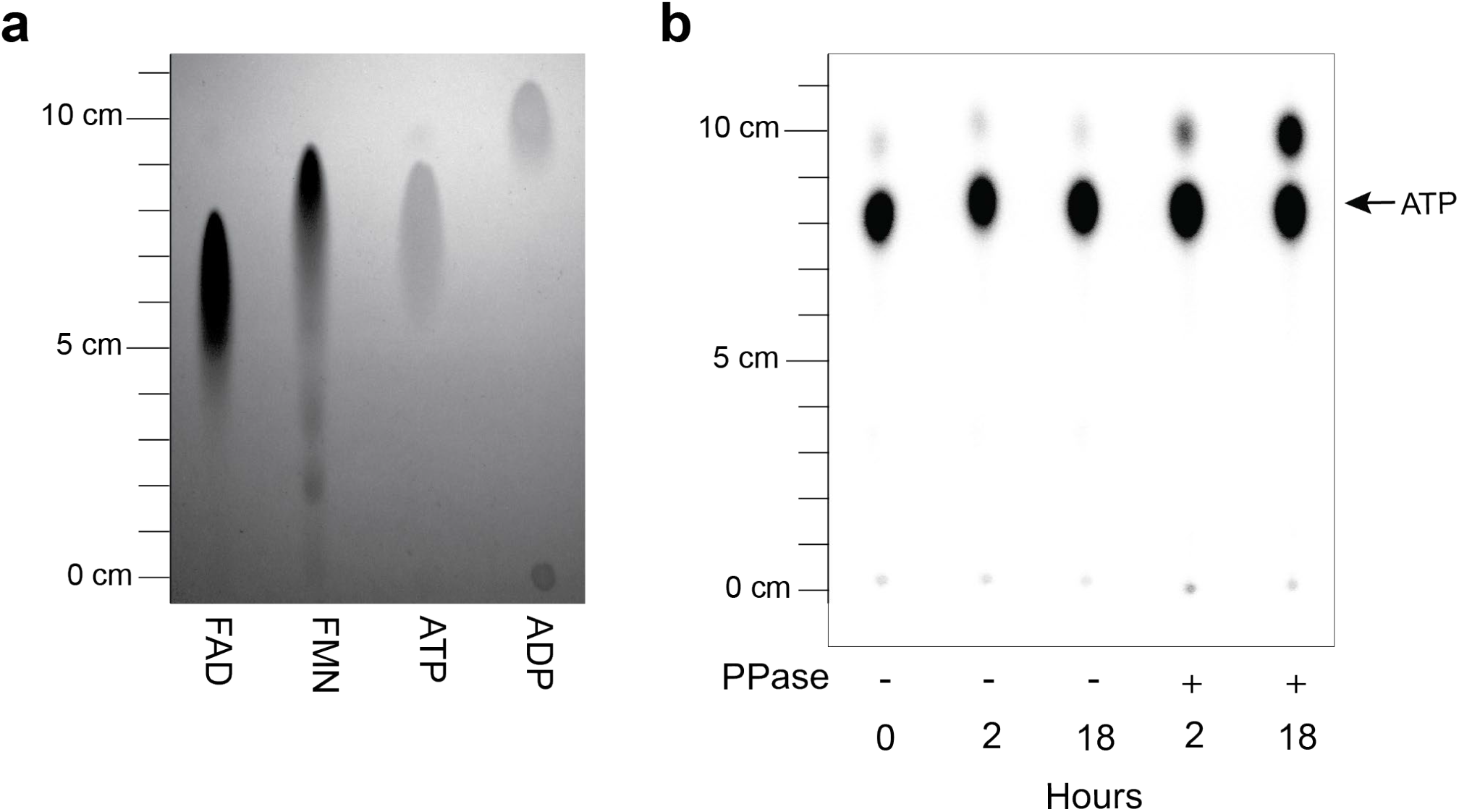
Attempted FADS enzymatic synthesis of ^32^P-FAD resulted in aberrant ^32^P product with PPase inclusion. (a) FMN, FAD, ATP and ADP standards were resolved with TLC using PEI cellulose F plates and imaged following excitation at 254 nm. (b) Attempted enzymatic reaction to synthesize radiolabeled ^32^P-FAD by adenylating FMN with ^32^P-ATP via recombinant FAD Synthetase. Reactions were setup as indicted in supporting methods, samples obtained at indicated time points and resolved with TLC, and imaged following exposure with a phosphorimaging screen. A ^32^P-labeled byproduct specifically appeared after incubation with PPase (lanes 4,5), with higher abundance at 18 hours (lane 5).

## Detailed Experimental Procedures

### Thin Layer Chromatography of Standards

#### Materials

##### Standards

ATP, A9062-5X1G, Millipore Sigma, >95%; ADP, A2754-5X1G, Millipore Sigma, >95%; AMP, A1752-5G, Millipore Sigma, >99%; NADH, 606-68-8, Avantor, >95%; NADP, 24292-60-2, Avantor, >95%; NAD^+^, N1636-25MG, Millipore Sigma, >99%; FMN, F6750-5G, Millipore Sigma, >73%; FAD, F6625-250MG, Millipore Sigma, >95%

TLC PEI Cellulose F plates: EMD Millipore, Cat#: 1057250001; Thin Layer Chromatography Tank; 254 nm UV Lamp

1. Mark TLC plate with a pencil at 2 cm above edge (origin), 7 cm above edge (5 cm resolve mark) and 17 cm above edge (15 cm resolve mark).
2. 5 μL of 50 μM standards of all nucleotides were spotted onto a TLC PEI Cellulose F plate at 2 cm above the bottom edge of the plate with 2.5 cm between each spot.
3. 50 mL of 18 MΩ H_2_O was added to a 30 cm x 27 cm x 10 cm glass chromatography tank such that the liquid was ∼1.5 cm deep.
4. The plate was resolved in the tank until the liquid reached 5 cm above origin. The plate was removed and allowed to dry on the bench top.
5. 50 mL of 1.4M LiCl was added to tank and the dry plate was placed back inside. The plate was resolved in the tank until the liquid reached 15 cm above origin.
6. The plate was allowed to dry on bench top and then imaged using a 254 nm UV lamp.

### NMNAT1 Synthesis of ^32^P-NAD^+^ from ^32^P-ATP

#### Materials

##### α-^32^P-ATP

Perkin Elmer Cat #: BLU003H250UC, Batches 07142, 03102, and 12091; Recombinant Human NMNAT1: (MyBioSource Cat#: MBS206313, Lot#: 21320702); Inorganic Pyrophosphatase (PPase): NEB Cat#: M0361S, Lot#: 10117903

1. The reaction mixture was assembled in 20 μL volume total, containing 3 μM α-^32^P-ATP, 50 μM ATP, 50 μM NMN and 5 μM Recombinant Human NMNAT1 in a final buffer of 50 mM Tris-HCl pH 7.4, 100 mM NaCl, 12 mM MgCl_2_, 1 mM DTT, and 0.1 units (1 μL) inorganic pyrophosphatase as indicated.
2. The reaction was carried out for 2 hours at room temperature, at which time 1 μL was removed and spotted onto a TLC PEI Cellulose F Plate and resolved as described in section “Thin Layer Chromatography of Standards”.
3. Upon completion, the reaction was snap-frozen in liquid nitrogen and stored at -20 °C.
4. After drying, the plate was wrapped in clingwrap and placed in a BioRad exposure cassette with a BioRad phosphorimaging screen for 3 minutes, then the screen was imaged on a Typhoon FLA 9500 Imager. The intensity of the dots was quantified using the ImageJ Gel Analyzer feature.

### Recovery of NAD^+^ from Cellulose TLC Plate

1. To recover NAD^+^ from cellulose plate, nucleotide spots were visualized with a 254 nm UV lamp and the NAD^+^ spot was circled with a pencil.
2. Cellulose in the circled area was removed with a razorblade and incubated with 500 μL of buffer (120 mM KCl, 5 mM KH_2_PO_4_, 1 mM EGTA, 3 mM HEPES-KOH pH 7.4) for 30 minutes at RT.
3. Buffer and cellulose were filtered through a 0.45 μm cellulose syringe filter to recover ^32^P-NAD^+^.

### Analysis and Quantification of Phosphorimaging Data

1. Import image into Image J. Use the rectangle tool and draw a rectangle around one lane to mark the ROI.
2. Use the copy function to duplicate the ROI for marking additional lanes. Repeat until all lanes are marked. Create plot profiles of each lane.
3. Use the Straight Line to draw a line across the background level on the plot profile to separate individual peaks.
4. Use the Magic Wand tool to highlight each peak. Magic Wand tool will automatically create area under the curve values for each peak selected.
5. Add up the values from each lane to sum the total intensity for that lane. Divide the intensity of any individual peak to get the percent of that species present, i.e. NAD^+^ peak over total sum to determine percent NAD^+^ of total.
6. Calculate the percent conversion of each species across replicate reactions. Using GraphPad, place replicates into a column data sheet. Compare reaction conditions with a Student’s t-test and calculate p-values.

### ^1^H-NMR of NMNAT1 Reaction with and without PPase

1. The reaction mixture was assembled in 100 μL volume total, containing 10 mM ATP, 10 mM NMN and 5 μM Recombinant Human NMNAT1 in a final buffer of 50 mM Tris-HCl pH 7.4, 100 mM NaCl, 12 mM MgCl_2_, 1 mM DTT, and 0.5 units (5 μL) inorganic pyrophosphatase as indicated.
2. Incubate mixture at room temperature for 16 hours.
3. Dilute reaction mixture 1:10 in Milli Q H_2_O to get 1 mM final concentration in 1 mL volume.
4. Add 10% D_2_O to the reaction and place in glass NMR tube.
5. Measure ^1^H-NMR using an Agilent 400 MHz NMR Magnet instrument.
6. ^1^H-NMR spectra analyzed using MestReNova 14.1 software.

### NAD^+^ Uptake into Isolated Yeast Mitochondria Expressing Human SLC25A51

#### Materials

##### Buffer A (no NAD^+^)

120 mM KCl, 5 mM KH_2_PO_4_, 1 mM EGTA, 3 mM HEPES-KOH pH 7.4

##### Buffer B (unlabeled NAD^+^ only)

120 mM KCl, 5 mM KH_2_PO_4_, 1 mM EGTA, 3 mM HEPES-KOH pH 7.4, 100 μM NAD^+^

##### Uptake Buffer (unlabeled NAD^+^ and ^32^P-tracer NAD^+^)

120 mM KCl, 5 mM KH_2_PO_4_, 1 mM EGTA, 3 mM HEPES-KOH pH 7.4, 100 μM NAD^+^, 2.9 nM ^32^P-NAD^+^

0.22 μm mixed-cellulose ester filter (Whatman, Cat#: WHA10401706)

KNF Vacuum Pump UN726

Wheaton 25 mm Filtration Assembly (Cat#: 419325)

MitoTracker Red CMXROS (ThermoFisher Cat #: M46752)

##### Yeast Strains used for isolation of fresh mitochondria

*Δndt1 Δndt2* BY4727 with PRS415-CEN/ARS-TEF-LEU plasmids expressing either Empty Vector (EV) or a yeast codon optimized human SLC25A51 (SLC25A51) sequence.

1. Yeast Mitochondrial Isolation was performed as previously described.^1-2^
2. In short, 1 L of yeast was grown in YP + 2% Raffinose ∼16 hours until reaching OD 600 of 3.0. Mitochondria were isolated using zymolyase, dounce homogenization and differential centrifugation to enrich for the mitochondrial fraction.
3. Mitochondria were kept on ice and used within 3 hours of isolation. Mitochondria were confirmed to still be functional using MitoTracker Red to show an intact membrane potential.
4. 1 μL of the reaction was diluted 1:20 (5 μL in 95 μL) into Buffer A to create a working stock of ^32^P-NAD^+^ (∼147 nM). This ^32^P-NAD^+^ stock was further diluted 1:50 (5 μL in 245 μL) into Buffer B to create Uptake Buffer. The final concentration of tracer ^32^P-NAD^+^ was ∼2.9 nM.
5. 1 mg of isolated mitochondria^1-2^ was resuspended in 50 μL Uptake Buffer and incubated for 5 min at room temperature. EV samples were included in parallel as paired negative controls.
6. At 5 minutes, the 50 μL reaction was stopped by adding 950 μL (20X volume) ice-cold Buffer B that only contained unlabeled NAD^+^.
7. The sample was filtered through a 0.22 μm mixed-cellulose ester filter using the Filtration Assembly attached to the vacuum pump. Filter was washed with an additional 5 mL of ice-cold Buffer A containing no NAD^+^.
8. The filter was covered in clingwrap and placed in a BioRad exposure cassette with a BioRad phosphorimaging screen for 16 hours, then the screen was imaged and analyzed as described in sections “NMNAT1 Synthesis of ^32^P-NAD^+^ from ^32^P-ATP” and “Analysis and Quantification of Phosphorimaging Data”.
9. Signal from EV negative control was averaged and the individual replicates from the EV and SLC25A51 conditions were normalized back to the paired EV average from that experiment.

### NAD Kinase Synthesis of ^32^P-NADP from ^32^P-NAD^+^

#### Materials

NADK (Adipogen Cat#:AG-40T-0091-C050, Lot#: A01419)

1. The reaction mixture was created in 20 μL volume with 0.44 μM ^32^P-NAD^+^, 100 μM ATP, 50 μM NAD^+^ and 5 μM NADK in a final buffer of 10 mM Tris-HCl pH 7.4, 100 mM NaCl, 5 mM MgCl_2_).
2. The reaction was carried out for 18 hours at room temperature. At 0.5, 2 and 18 hours, 1 μL was removed and spotted onto a TLC PEI Cellulose F Plate at 2 cm above the bottom edge, with 2.5 cm between each spot, and resolved as described above.
3. After drying, the plate was wrapped in clingwrap and placed in a BioRad exposure cassette with a BioRad phosphorimaging screen for 3 minutes, then the screen was imaged and analyzed as described in sections “NMNAT1 Synthesis of ^32^P-NAD^+^ from ^32^P-ATP” and “Analysis and Quantification of Phosphorimaging Data”.

### Attempted FAD Synthetase Reaction from ^32^P-ATP

#### Materials

Recombinant human FADS (MyBioSource Cat#: MBS1342413, Lot#: YA05272b1g5)

1. The reaction mixture was assembled in 20 μL volume with 3 μM α-^32^P-ATP, 50 μM ATP, 50 μM FMN and 5 μM human FADS in a final buffer of 50 mM Tris-HCl pH 7.4, 100 mM NaCl, 12 mM MgCl_2_ and 1 mM DTT^3^, with the addition of 0.1 Units (1 μL) inorganic pyrophosphatase in the proper conditions.
2. The reaction was carried out for 18 hours at room temperature. At 2 and 18 hours, 1 μL was removed and spotted onto a TLC PEI Cellulose F Plate at 2 cm above the bottom edge, with 2.5 cm between each spot, and resolved as described above.
3. After drying, the plate was wrapped in clingwrap and placed in a BioRad exposure cassette with a BioRad phosphorimaging screen for 3 minutes, then the screen was imaged and analyzed as described in sections “NMNAT1 Synthesis of ^32^P-NAD^+^ from ^32^P-ATP” and “Analysis and Quantification of Phosphorimaging Data”.

## Notes

Supporting Information Placeholder

### Competing Interest Statement

The authors have declared no competing interest.

